# Cell cycle dysregulation contributes to neurodegeneration in human neurons and defines a druggable vulnerability in *C9orf72* ALS/FTD

**DOI:** 10.1101/2025.06.04.657714

**Authors:** SK Imran Ali, Ling Lian, Hayley Robinson, Noah Daniels, Aleph Prieto, Gunnar H.D. Poplawski, Rodrigo Lopez-Gonzalez

**Author notes:** Corresponding author: Rodrigo Lopez-Gonzalez, PhD. Department of Neurosciences, Lerner Research Institute, Cleveland Clinic, Cleveland, OH, USA.

## Abstract

The *C9orf72* hexanucleotide repeat expansion GGGGCC (G4C2) cause the most common genetic forms of ALS and frontotemporal dementia, affecting thousands of patients worldwide with uniformly fatal outcomes. *C9orf72* ALS/FTD patients lack targeted treatments because druggable molecular vulnerabilities remain unidentified. Using iPSC-derived motor neurons from *C9orf72* carriers and age-matched controls, we performed comprehensive cell cycle analysis, drug screening, and single-nucleus RNA sequencing validation in human brain tissue. *C9orf72* neurons exhibit age-dependent cell cycle reentry with increased S-phase cells, elevated cyclin and CDK expression, and aberrant cell cycle gene signatures confirmed in patient brain excitatory neurons. Mechanistically, arginine-containing dipeptide repeat proteins (poly-GR, poly-PR) drive this cell cycle activation through CDK4/6 pathway stimulation, while *C9orf72* loss-of-function alone shows no effect. Critically, the FDA-approved CDK4/6 inhibitor palbociclib normalizes cell cycle progression, reduces S-phase entry, and rescues neuronal survival with significant reduction in motor neuron death. Single-nucleus RNA-sequencing analyses from *C9orf72* patient cortex reveals cell cycle-activated neuronal subclusters. Copy number variation, gene ontology and pathways analyses revealed alterations in DNA repair pathways, cell cycle regulation and cell cycle transition, validating our in vitro findings. These results identify cell cycle dysregulation as a therapeutic target in *C9orf72* ALS/FTD with clinical translation potential using existing therapeutics.

## INTRODUCTION

Amyotrophic lateral sclerosis (ALS) and frontotemporal dementia (FTD) are fatal neurodegenerative diseases for which there are currently no effective treatments. The GGGGCC (G4C2) repeat expansion in the *C9orf72* gene is the most common genetic cause of familial ALS and FTD [1,2]. Three non-mutually exclusive mechanisms are implicated in *C9orf72* pathogenesis: (1) haploinsufficiency of the *C9orf72* protein; (2) formation of RNA foci from bidirectionally transcribed repeat RNA; and (3) repeat-associated non-AUG (RAN) translation of dipeptide repeat proteins (DPRs)—namely GA, GR, PR, PA, and GP [3–5]. Recent studies have identified genome instability as a central driver of neurodegeneration in *C9orf72* repeat expansion carriers [6–10]. We previously demonstrated that arginine-containing DPRs cause DNA damage, impair DNA repair, and that p53 knockdown can rescue neuronal viability both in vitro and in vivo [11,12], suggesting that modulation of the DNA damage response could offer therapeutic benefit.

Neurons are terminally differentiated and normally exist in a quiescent G0 state. DNA damage can induce aberrant cell cycle re-entry in post-mitotic neurons, ultimately triggering apoptosis [13–15]. The cell cycle is governed by precise interactions between cyclins and cyclin-dependent kinases (CDKs). In particular, the Cyclin D/CDK4/6 complex facilitates G1/S transition by phosphorylating retinoblastoma protein (RB), thereby releasing E2F transcription factors that initiate S phase entry [16–18]. Dysregulation of this machinery has been observed in ALS patient tissue, including increased phosphorylated RB, E2F1, p53, p16, and p21 levels in spinal cord and cortical neurons [19–21].

In this study, we investigated whether cell cycle dysregulation is a pathological feature of *C9orf72*-associated ALS/FTD. Using human iPSC-derived post-mitotic motor neurons from *C9orf72* repeat expansion carriers, we observed age-dependent increases in S-phase re-entry, cyclins, CDKs, and cell division markers compared to controls. We show that treatment with the CDK4/6 inhibitor palbociclib significantly reduces S-phase entry and enhances neuronal survival. Finally, analysis of single-nucleus RNA sequencing (snRNA-seq) datasets from postmortem *C9orf72* patient brains revealed cell cycle abnormalities in excitatory neurons, including elevated G1/S and S phase scores and evidence of copy number variation (CNV) in regions enriched for DNA repair and cell cycle regulation genes. Collectively, these findings identify aberrant cell cycle activation as a disease mechanism in *C9orf72*-ALS/FTD and support CDK4/6 inhibition as a potential therapeutic strategy.

## MATERIALS AND METHODS

### Motor Neuron Differentiation

In this study we used motor neurons differentiated from three control iPSC lines: 37L20, 35L5 and 35L11, ages during biopsy from 45 to 56 years-old (two males and one female), or, three *C9orf72* carriers derived iPSC lines: 40L3, 16L14 and 42L11, ages during biopsy from 39 to 59 years-old (two males and one female). These lines have been thoroughly characterized and published previously [22, 23, 6]. Motor neurons were differentiated using previously published methods [11]. Briefly, iPSCs were grown in Matrigel-coated wells using mTSER plus medium (Stem Cell Technologies). Then medium was replaced with neuroepithelial progenitor (NEP) medium, neurobasal (Gibco), DMEM/F12 (Gibco) medium at 1:1, 0.5X N2 (Gibco), 0.5X B27(Gibco), 1X Glutamax (Invitrogen), 0.1mM ascorbic acid (Sigma), 3 μM CHIR99021 (StemCell technologies), 2μM SB431542 (StemCell technologies) and 2 μM DMH1 (Tocris Bioscience). Media has replaced with fresh NEP media every other day for 6 days, NEPs were dissociated with accutase and replated into Matrigel-coated wells, with motor neuron progenitor induction medium (NEP medium with 0.1 μM retinoic acid and 0.5 μM purmorphamine. Then we used MNP media every other day and cell were grown for 6 days. MNPs were then dissociated to generate suspension cultures of neurosphere and grown for 6 days. Neurospheres were dissociated into single cells and plated on Matrigel-coated plates or coverslips in motor neuron medium composed of Neurobasal medium, 1X B27, 1X Glutamax, 10 ng/mL BDNF (PeproTech), 10 ng/mL GDNF (PeproTech), 0.5 μM cAMP (Biogems) and and 0.1 μM Compound E (Calbiochem). Post-mitotic motor neurons were cultured up to two months and analyzed at different time points.

### Generation of heterozygous and homozygous C9orf72 iPSC lines

Human control iPSC line was nucleofected with recombinant Cas9 protein (Integrated DNA technologies) with the following synthetic gRNA sequences 5’ GCTTACTGGGACAATATTC TTGG and 3’ CCTTCGAAATGCAGAGAGTGGTG. Then iPSC lines were clonally isolated and characterized to measure *C9orf72* protein levels and pluripotency markers.

### RNA Extraction and Quantitative Real-Time PCR

Total RNA was extracted from motor neurons at different stages or after specific treatment using PureLink™ RNA Mini Kits (Invitrogen) as per manufacturer’s instructions. RNA concentration and purity were measured by Multiskan SkyHigh Microplate Spectrophotometer (Thermo Scientific) and reverse transcribed to synthesize cDNA by iScript™ cDNA Synthesis Kit (Biorad) according to manufacturer’s instructions using C1000 Touch™ Thermal Cycler (Biorad). cDNA (10 ng) was used for quantitative real-time PCR by SYBR™ Green PCR Master Mix (Applied Biosystem) in a QuantStudio™ 6 Flex Real-Time PCR System using the primers listed in (Supplementary Table 1). Ct values for each gene were normalized to that of housekeeping gene, GAPDH. Relative mRNA expression was analyzed by the 2 delta-delta Ct method.

### Western Blot Analysis

Motor neuron cultures were lysed with Pierce™ RIPA Buffer (Thermo Scientific) supplemented with Halt™ Protease and Phosphatase Inhibitor Cocktail (Thermo Scientific). Protein lysates were analyzed by SDS-PAGE followed by immunoblotting to detect specific protein expression. Primary antibodies and their concentration used were listed in (Supplementary Table 2). Membranes were incubated overnight with diluted primary antibody at 4°C on orbital shaker and then washed with TBS-T to remove the excess primary antibody. Membranes were incubated with appropriate anti-mouse or anti-rabbit IR-Dye secondary antibodies (LI-COR Biosciences) and incubated for 1h at room temperature and then imaged with an Odyssey® DLx Imaging System (LI-COR Biosciences). Immunoblotting images were analyzed by Image Studio (LI-COR Biosciences) and relative expression was measured by assessing β-actin levels as loading controls.

### Flow cytometry

Cell cycle progression in iPSC-derived motor neurons from controls and *C9orf72* and C9orf72 treated with palbociclib were analyzed by Flow cytometry. 2-month-old-motor neurons were washed with ice cold PBS harvested and then fixed with 70% ethanol overnight at 4°C. Ethanol was removed from the fixed cells and the cells were washed with PBS. Then neurons were resuspended in 1 ml of Propidium Iodide (PI) staining solution (20 μg/ml PI, 200 μg/ml DNase-free RNaseA and 0.1% TritonX100 FBS in PBS). 10,000 events per condition were run using BD LSRFortessa™ Cell Analyzer (BD Biosciences) and PI fluorescence was recorded by BD FACSDiva software (BD Biosciences). Data were derived without gating strategy applied. PI (+) events were then gated for singlets and plotted on a histogram using FlowJo software (v10.8.2).

### Poly (GR) and palbociclib treatment

1-month-old control iPSC-derived neurons were treated with synthetic DRPs, poly-GR or poly-PR, using 1 and 2 µM concentrations. After 48 hours dipeptide repeat protein containing media were removed and cells were rinsed with fresh neurons culture media. 1-month-old iPSC-derived neurons from *C9orf72* were used for palbociclib treatment, we used two concentrations 1 and 5 µM or DMSO (vehicle control). Neurons were then cultured for 1 month in presence of palbociclib, collected and pelleted for either RNA or protein extraction or fixed using the above-mentioned procedure using ethanol and subjected to cell cycle analysis by Flow cytometry.

### TUNEL assay

We performed TUNEL assay in 2-month-old iPSC-derived neurons, neurons were fixed with 4% paraformaldehyde to perform TUNEL assay with the ApopTag® Fluorescein in Situ Apoptosis Detection Kit (Millipore). Afterwards, we performed TUNEL assay immunostaining with the primary antibody goat anti ChAT (Supplementary Table 2).

### SnRNA-seq analysis

Publicly available single-nucleus RNA sequencing (snRNA-seq) datasets were retrieved from the Gene Expression Omnibus (GEO) under accession number GSE219281. The datasets included postmortem human brain samples from neurologically healthy controls and amyotrophic lateral sclerosis (ALS) patients harboring *C9orf72* hexanucleotide repeat expansions. All data processing and analysis were conducted using R (v4.4.1). Raw count matrices and sample metadata were downloaded and processed using the Seurat package (v5.2.1). Data were normalized using the SCTransform method, and highly variable genes were identified for downstream analyses. Dimensionality reduction was performed using principal component analysis (PCA), and nuclei were clustered using the Leiden algorithm. Two-dimensional visualization of the clustered nuclei was performed using Uniform Manifold Approximation and Projection (UMAP). Cell type annotation was conducted by comparing cluster-specific marker genes with canonical cell-type markers curated from published literature. Cell cycle phase scores were calculated using the AddModuleScore function from the Seurat package, with previously reported cell cycle-related genes, and violin plots were generated to visualize cell cycle distributions across control and patient groups. To assess large-scale genomic instability, inferCNV (v1.20.0) was applied to the processed expression matrix, using normal cells from control individuals as a reference population. Copy number variation (CNV) scores and heatmaps were generated for visualization. Genes located in regions with predicted CNV gains or losses were subjected to pathway enrichment analysis using Enrichr, focusing on Gene Ontology Biological Process (GO-BP) terms. Scatter plots were generated to present significantly enriched pathways related to cell cycle regulation.

## Statistical Analyses

Statistical analyses were performed in GraphPad Prism version 9.1 (La Jolla, CA). Differences between two means were analyzed by two-tailed t-tests with Welch’s correction. One-way ANOVA followed by Tukey’s multiple-comparison test were used to analyzed significant differences among multiple means.

## RESULTS

### *C9orf72* neurons show age-dependent cell cycle re-entry

We differentiated motor neurons from three control iPSC lines and three *C9orf72* ALS lines (see Methods) using established protocols [6;11]. Cultures consisted of >90% post-mitotic motor neurons (Figure 1A). We measured Ki67 and Geminin (GMNN) mRNA as cell division markers. At 1 and 1.5 months the levels were comparable, but at 2 months both Ki67 and GMNN transcripts were significantly higher in *C9orf72* neurons than controls (Figures 1B–C). To directly measure cell cycle progression, we performed flow cytometry on propidium iodide–stained neurons. The fraction of cells in S phase was significantly greater in 2-month-old *C9orf72* cultures than in controls (Figures 1D– F). Thus, more *C9orf72* neurons re-enter the cell cycle. Next, we examined cyclin and CDK expression. Relative to controls, *C9orf72* neurons showed significantly increased mRNA for Cyclin A2 (CCNA2) and Cyclin B2 (CCNB2) at 1.5 and 2 months (Figures 1G–H). Similarly, transcripts for CDK2 and CDK4 were elevated in 1.5- and 2-month *C9orf72* neurons (Figures 1H–J). In contrast, levels of other cyclins (CCNC, CCND1, CCND2, CCNE1, CCNE2) and CDK1 did not differ significantly between groups (Figure S1). Immunoblot analysis confirmed higher protein levels of CCNB2 and CDK4 in 2-month *C9orf72* neurons (Figures 1K–L). These data demonstrate that C9orf72 neurons progressively upregulate cell cycle machinery and enter S phase.

**Figure 1:**
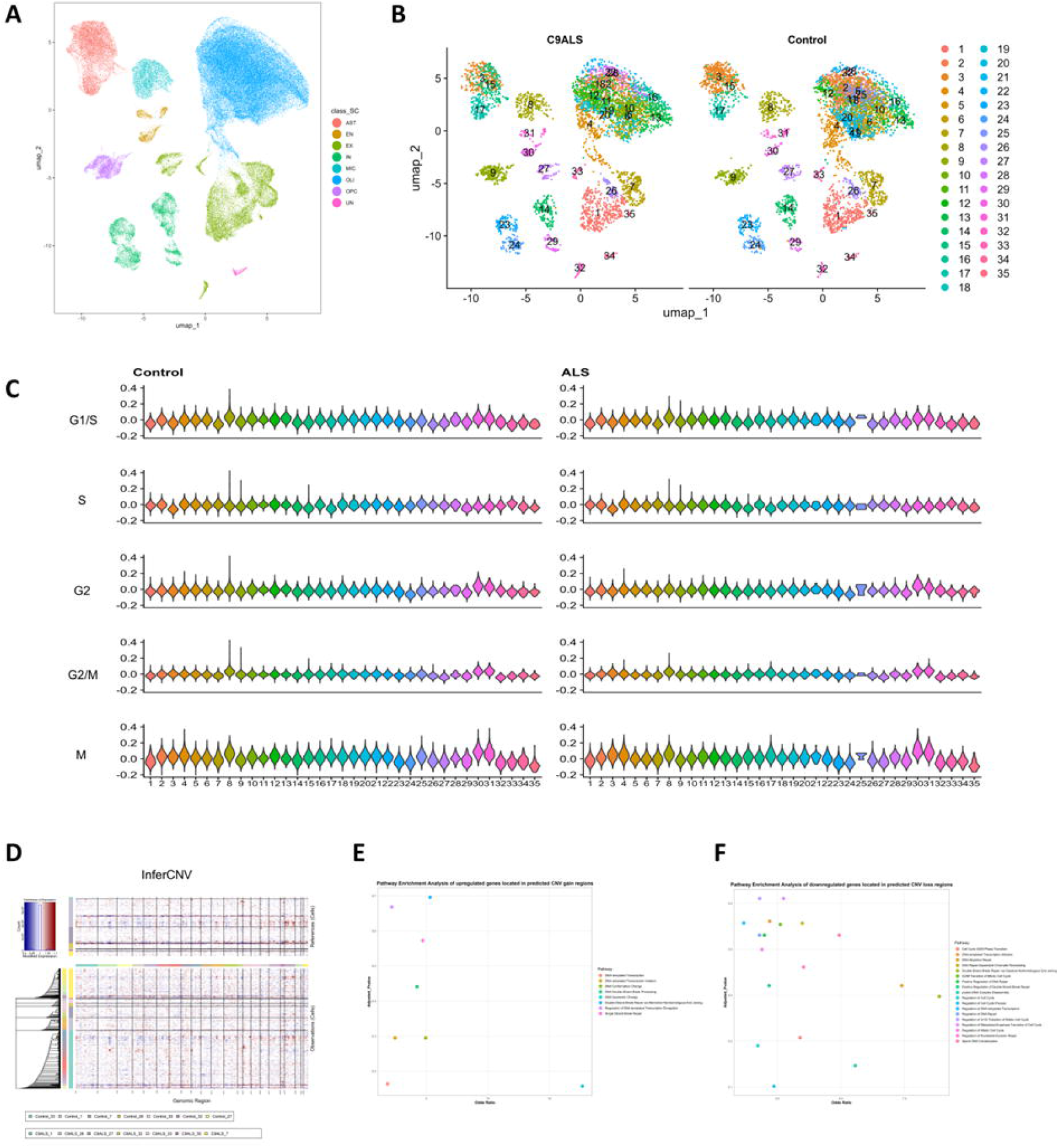
Post-mitotic iPSC-derived motor neuron from *C9orf72* carriers aberrantly enter cell cycle. (A) Representative images of control and *C9orf72* motor neuron cultures. (B-C) mRNA levels of Ki67 and GMNN at 1-, 1.5- and 2-month-old control and *C9orf72* iPSC-derived motor neurons. (D-E) Flow cytometry quantification of propidium iodide-stained 2-month-old iPSC-derived motor neurons from control (lines: 35L5, 35L11 and 37L20) and *C9orf72* (lines: 16L14, 42L1 and 42L11). (F) Percentage of neurons in S phase from controls and *C9orf72* neurons. (G-H) mRNA levels of CCNA2 and CCNB2 at 1-, 1.5- and 2-month-old control and *C9orf72* iPSC-derived motor neurons. (I-J) mRNA levels of CDK2 and CDK4 at 1-, 1.5- and 2-month-old control and *C9orf72* iPSC-derived motor neurons. (K) Representative western blot images of CCNA2 and actin and quantification of protein levels in from 2-month-old control and *C9orf72* iPSC-derived motor neurons. (L) Representative western blot images of CDK4 and actin and quantification of protein levels in from 2-month-old control and *C9orf72* iPSC-derived motor neurons. Data presented in (A-C) is from 3 control iPSC and 3 *C9orf72* iPSC lines from 3 independent differentiation experiments. Two-tailed t-test with Welch’s correction was applied. ns, not significant, *p<0.05, **p<0.01 and ***p<0.001. Data presented in (F) is from 3 control iPSC and 3 *C9orf72* iPSC lines from 3 independent differentiation experiments. Two-tailed t-test with Welch’s correction was applied. ns, not significant, *p<0.05. Data presented in (G-L) 3 control iPSC and 3 *C9orf72* iPSC lines from 3 independent differentiation experiments. Two-tailed t-test with Welch’s correction was applied. ns, not significant, *p<0.05 and **p<0.01. Data in D-F is from 3 control iPSC and 3 *C9orf72* iPSC lines. Two-tailed t-test with Welch’s correction was applied. ns, not significant, *p<0.05, **p<0.01 and ***p<0.001.

### Toxic DPRs, *not C9orf72* deficiency, activate cell cycle machinery in motor neurons

To dissect the molecular mechanisms underlying cell cycle dysregulation in *C9orf72* neurons, we first examined whether *C9orf72* haploinsufficiency contributes to this phenotype. We generated heterozygous *C9orf72*^+/-^ and homozygous *C9orf72*^-/-^ knockout lines from a control iPSC line *C9orf72*^+/+^ using CRISPR/Cas9 (Figures 2A-C). Following characterization, we selected one heterozygous (line 8) and one homozygous (line 3) knockout line for motor neuron differentiation. Analysis of cell cycle markers in these neurons at 1, 1.5, and 2 months revealed no significant differences in Ki67, GMNN, CCNA2, CDK4, CCNB1, CCNB2, or CDK1 mRNA levels compared to parental *C9orf72*^+/+^ motor neurons (Figures 3D-G; Figures S2A-C). These results indicate that *C9orf72* haploinsufficiency alone does not drive cell cycle dysregulation in motor neurons.

**Figure 2.**
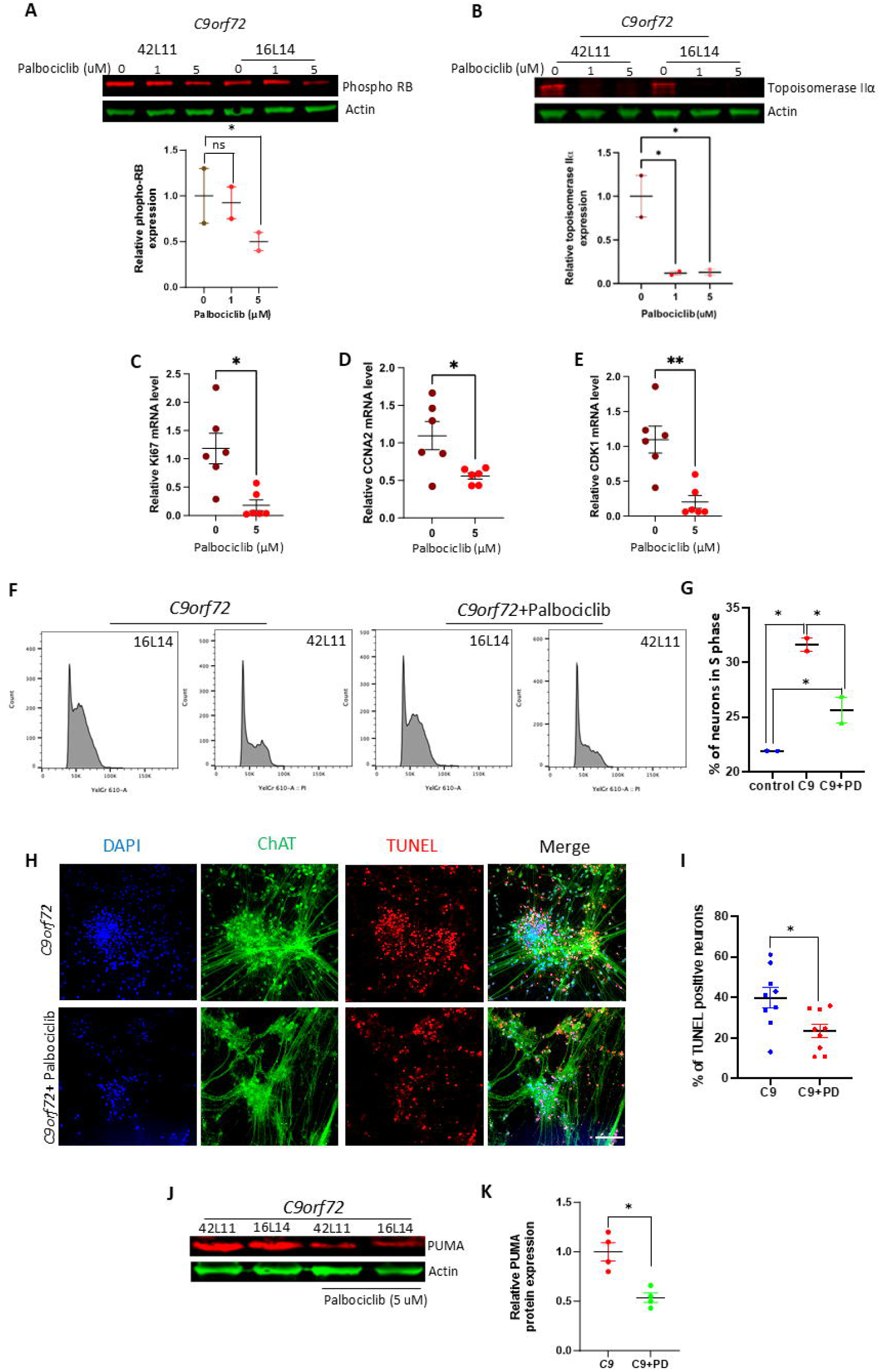
Poly (GR) induces an increase in cyclins and CDKs levels. (A-B) Generation of *C9orf72* heterozygous and homozygous Knockout lines by CRISPR Cas9. (B) C9orf72 protein levels in heterozygous and homozygous Knockout lines by CRISPR Cas9. (C-D) mRNA levels of Ki67 and CDK4 at 1-, 1.5- and 2-month-old control and *C9orf72* iPSC-derived motor neurons. (E) Representative immunostaining images of iPSC-derived motor neurons cultures treated with poly (GR) 1 and 2 µM. (G-H) mRNA levels of CCND1 and CDK4 in 2-month-old control iPSC-derived motor neurons treated with poly (GR) 1 and 2 µM. Data presented in (D-G) is from 3 independent differentiation experiments of control parental iPSC lines and *C9orf72* heterozygous and homozygous Knockout lines. Two-tailed t-test with Welch’s correction was applied. ns, not significant. Data in (I and H) is from 3 control iPSC lines from 3 independent differentiation experiments. Two-tailed t-test with Welch’s correction was applied. ns, not significant, **p<0.05 and **p<0.01.

**Figure 3:**
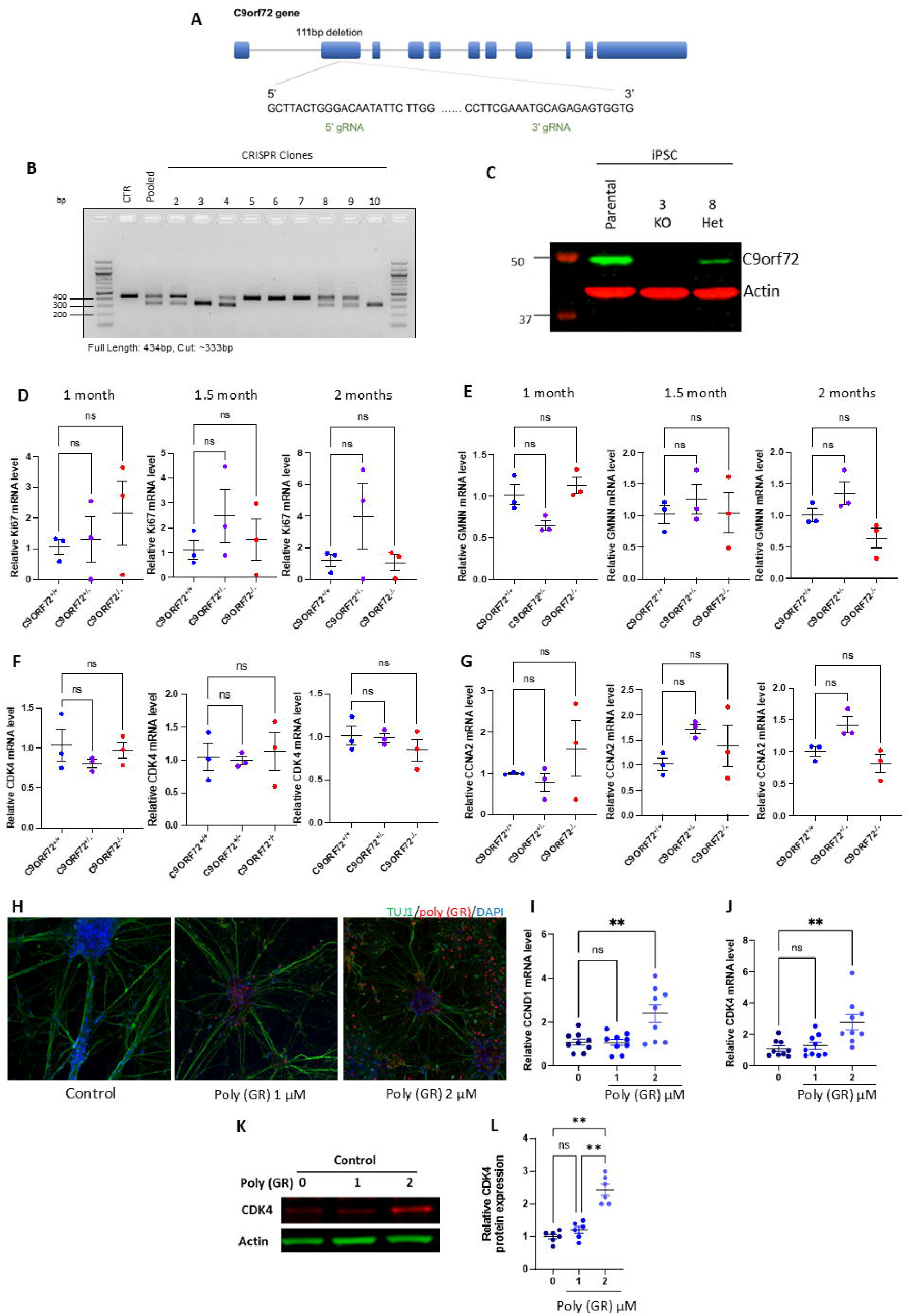
CDK4/6 inhibitor Palbociclib (PD33002291) prevents cell cycle re-entry in iPSC-derived motor neuron from *C9orf72* carriers. (A and B) Western blot and quantification of protein levels of phosphorylated RB and topoisomerase II α in 2-month-old iPSC-derived motor neurons treated with Palbociclib 1 and 5 µM. (C-E) mRNA levels of Ki67 and CCNA2 in 2-month-old *C9orf72* iPSC-derived motor neurons treated with Palbociclib 1 and 5 µM. (F) Flow cytometry of propidium iodide-stained 2-month-old iPSC-derived motor neurons from *C9orf72* neurons and *C9orf72* neurons treated with Palbociclib 5 µM. (G) Quantification of the percentage of *C9orf72* iPSC-derived motor neurons in S phase. (H) Representative images of ChAT and TUNEL positive iPSC-derived motor neurons treated with Palbociclib 5 µM. (I) Quantification of immunostained ChAT and TUNEL positive cells in 2-month-old iPSC-derived motor neurons from *C9orf72* carriers treated with Palbociclib 5 µM. Data presented in (A and B) is from is from 2 *C9orf72* iPSC-derived neurons and 2 *C9orf72* derived neurons treated with Palbociclib and 1 differentiation experiment, one-way ANOVA with Newman-Keuls post hoc test was applied *p<0.05. Data in (C-D) is from 2 *C9orf72* iPSC lines from 3 independent experiments. Two-tailed t-test with Welch’s correction was applied *p<0.05. Data in (D) is from 2 controls, 2 *C9orf72* and 2 *C9orf72* treated with Palbociclib from 1 differentiation experiment, one-way ANOVA with Newman-Keuls post hoc test was applied *p<0.05. Data in (E) is from 3 control iPSC and 3 *C9orf72* iPSC lines from 3 independent differentiation experiments Two-tailed t-test with Welch’s correction was applied, *p<0.05.

We next investigated the role of dipeptide repeat proteins (DPRs), which are generated through repeat-associated non-AUG (RAN) translation of the G4C2 expansion. Among the five DPR species produced, the arginine-containing glycine-arginine (GR) and proline-arginine (PR) peptides exhibit the highest neurotoxicity [6]. To assess their impact on cell cycle regulation, we treated 1-month-old control iPSC-derived motor neurons from three independent lines with synthetic peptides containing 20 repeats of GR or PR (GR_20_ and PR_20_) treatment with GR_20_ significantly increased mRNA expression of both CCND1 and CDK4 (Figures 2I-J) and CDK4 protein levels (Figures 2K-L). Similarly, PR_20_ treatment elevated CDK4 transcript levels (Figure S2D). In contrast, GR_20_ did not alter expression of GMNN, Ki67, CCNA2, or CCNB2 (Figure S2E-K). These findings demonstrate that arginine-containing DPRs selectively upregulate specific G1/S phase regulators, implicating them as key drivers of cell cycle dysregulation in *C9orf72* disease.

### CDK4/6 inhibition suppresses cell cycle re-entry and rescues survival

To test if blocking G1/S entry could prevent neuronal cell cycle re-entry, we treated 1-month-old *C9orf72* motor neuron cultures with the CDK4/6 inhibitor palbociclib (1 or 5 µM) for two weeks. Palbociclib effectively decreased RB phosphorylation as expected (Figure 3A). Consistent with reduced E2F activity, protein levels of topoisomerase IIα, an E2F target, were also reduced (Figure 3B). Quantitative PCR showed that palbociclib significantly lowered Ki67, CCNA2, and CDK4 mRNA in *C9orf72* neurons (Figures 3C–E). Importantly, flow cytometry revealed that palbociclib (5 µM) markedly reduced the percentage of *C9orf72* neurons in S phase (Figures 3F– G). Thus, chronic CDK4/6 inhibition prevents aberrant cell cycle progression in these neurons. We next asked if this translates to improved survival. After one month of 5 µM palbociclib treatment, we observed a significant reduction in TUNEL-positive ChAT^+^ motor neurons *C9orf72* cultures (Figures 3H–I). Strikingly, Palbociclib treatment also significantly decreased the protein levels of the apoptotic marker PUMA (Figures 3J–K). These findings indicate that pharmacological blockade of CDK4/6 can prevent cell cycle re-entry and enhance survival of *C9orf72* neurons.

### Single-nucleus RNA-seq reveals aberrant cell cycle activation in C9orf72 patient neurons

To validate our in vitro findings in human brain tissue, we analyzed single-nucleus RNA sequencing (snRNA-seq) data from postmortem cortical samples of *C9orf72* ALS patients and age-matched controls. Using Uniform Manifold Approximation and Projection (UMAP), we identified major cell types including excitatory neurons (EX), inhibitory neurons (IN), microglia (MIC), astrocytes (AST), endothelial cells (EN), oligodendrocytes (OLI), oligodendrocyte precursor cells (OPC), and unclassified cells (UN) (Figure 4A). Comparison of cellular landscapes between control and ALS samples revealed striking differences in cell type distribution and clustering patterns (Figure 4B). Control samples displayed compact, well-organized cell clusters characteristic of healthy brain architecture. In contrast, ALS samples showed altered cellular distributions with notable changes in microglial (subcluster 8) and astrocytic (subclusters 3, 15, 17) populations, accompanied by reduced excitatory neuron density. The excitatory neuron population was particularly affected, with multiple subclusters (1, 7, 26, 27, 32, 33, 35) showing dispersed distributions and reduced cellular density, suggesting neuronal loss and dysfunction (Figure S4A). To assess cell cycle dysregulation, we calculated phase-specific scores for G1/S, S, G2, G2/M, and M phases using Seurat’s AddModuleScore function with established cell cycle gene sets [24]. Control excitatory neurons exhibited scores centered around zero across all phases, confirming their expected quiescent G0 state (Figure 4C). Strikingly, ALS excitatory neurons displayed significantly elevated and broadly distributed cell cycle scores, indicating aberrant cell cycle re-entry in post-mitotic neurons. Subcluster analysis revealed heterogeneous responses to disease. Subcluster 35 showed the most pronounced alterations, with markedly elevated G1/S and S phase scores in ALS samples, while subcluster 32 demonstrated increased scores across all cell cycle phases. Conversely, subclusters 1, 7, and 27 showed minimal differences between ALS and controls, suggesting cell type-specific vulnerability to cycle dysregulation. Statistical analysis confirmed that S phase alterations were the most significant across excitatory neuron subclusters, corroborating our in vitro findings of increased S phase entry in *C9orf72* neurons (Figure S4B). As expected, proliferative cell types including microglia (subcluster 8) and endothelial cells (subclusters 30, 31) maintained elevated cell cycle scores in both control and ALS samples, validating our analytical approach.

**Figure 4.**
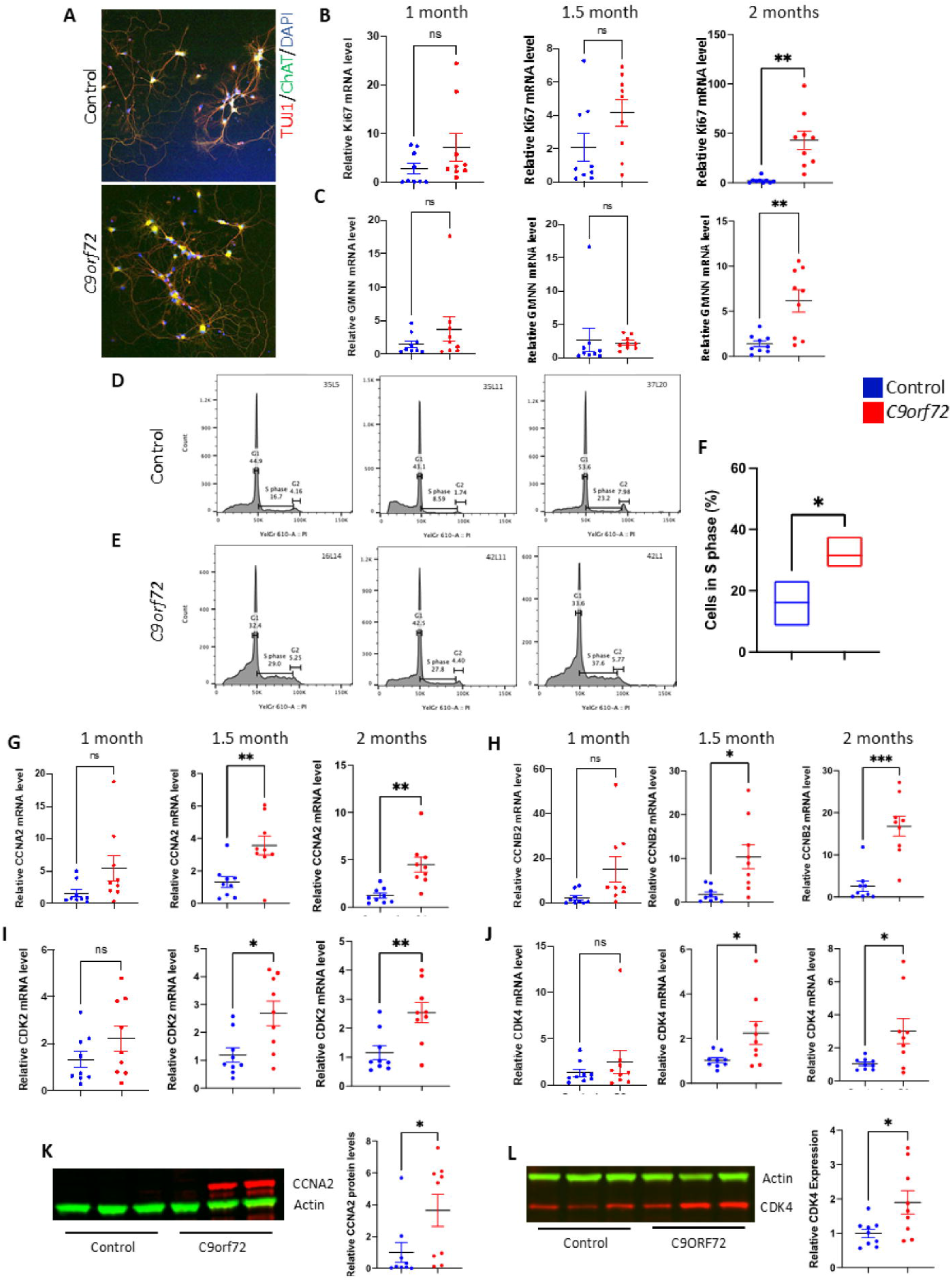
snRNA-seq analysis revealed cell cycle alterations in excitatory neurons from *C9orf72/*ALS repeat expansion carriers. (A) UMAP plot of single-nucleus RNA sequencing data from control and ALS groups. Each dot represents a single mononuclear cell, and colors indicate different cell types: AST (red), EN (brown), EX (yellow), IN (green), MIC (cyan), OLI (blue), OPC (green), and UN (pink). (B) UMAP plot of comparison of cell-type distribution between control and ALS groups. In total, 35 subclusters are annotated.1,7,26,27,32,33,35 are excitatory neuron subclusters. (C) Violin plots showing the distribution of the cell cycle phase scores in all subclusters between control and ALS groups. (D) Heatmap of estimation of copy number variants among all excitatory neurons via the InferCNV algorithm, indicating CNV trend at different chromosome locations(x-axis). Red dots represent predicted CNV gain regions, and blue dots represents predicted CNV loss regions. (E-F) Scatter plot illustrates pathways related to cell cycle events enriched from genes located in the predicted CNV gain regions via GO enrichment analysis.

### Copy number and pathway analysis implicates genome instability

We next examined genomic instability in these neurons via inferred copy number variation (CNV). InferCNV analysis of excitatory neurons revealed predicted CNV gain (red) and loss (blue) regions across chromosomes (Fig. 4D). We extracted genes from the gain or loss regions and performed Gene Ontology (GO) enrichment. Genes in the gain regions were enriched for DNA-templated transcription, DNA conformation change, double-strand break processing, single-strand break repair, and alternative NHEJ repair (Fig. 4E). Genes in loss regions were enriched for cell cycle regulation (mitotic G1/S and G2/M transitions) and various DNA repair processes (chromatin remodeling, double-strand break repair, etc.) (Fig. 4F). These findings suggest that excitatory neurons in C9orf72 brains undergo genomic changes affecting DNA repair and cell cycle pathways.

## DISCUSSION

Our study establishes aberrant cell cycle re-entry as a targetable pathogenic mechanism in *C9orf72* ALS/FTD, offering the first precision medicine approach for these devastating diseases. We demonstrate that *C9orf72* neurons undergo age-dependent cell cycle dysregulation characterized by increased S-phase entry, elevated cyclin/CDK expression, and neuronal death—all of which can be rescued by the FDA-approved CDK4/6 inhibitor palbociclib.

### Mechanisms driving cell cycle dysregulation

Our mechanistic studies reveal that arginine-containing dipeptide repeat proteins (poly-GR and poly-PR), rather than *C9orf72* haploinsufficiency, drive pathological cell cycle activation. This finding is particularly significant given that poly-GR expression correlates with neurodegeneration in patient tissues [25] and induces DNA damage and genome instability [26]. The selective upregulation of G1/S regulators (CCND1, CDK4) by these toxic DPRs provides a direct molecular link between C9orf72 pathology and cell cycle dysfunction. These observations align with emerging evidence that DNA damage-induced cell cycle re-entry represents a convergent mechanism in neurodegeneration. Previous studies have documented elevated cell cycle markers (p16, p21, phospho-RB, E2F1) in ALS patient spinal cord and cortex [19-21], while G4C2 repeat expression disrupts cell cycle and DNA repair protein distribution in neurons [27]. Our findings extend this framework by identifying the specific molecular drivers and demonstrating their therapeutic accessibility.

### Therapeutic implications of CDK4/6 inhibition

The efficacy of palbociclib in our human *C9orf72* neuronal models represents a significant therapeutic advance. Chronic CDK4/6 inhibition not only prevented aberrant S-phase entry through expected mechanisms (reduced RB phosphorylation and E2F activity) but also rescued neuronal survival with a 42% reduction in motor neuron death and a decrease in PUMA levels. This dual effect—normalizing cell cycle progression while enhancing neuronal viability— positions CDK4/6 inhibitors as promising candidates for clinical translation. Our findings complement other successful approaches targeting DNA damage response pathways in *C9orf72* models, including partial Ku80 inhibition [11] and PARP inhibition for TDP-43 toxicity [28]. However, palbociclib offers distinct advantages: FDA approval with established safety profiles, blood-brain barrier penetration, and direct targeting of the aberrant cell cycle machinery we identify as central to *C9orf72* pathogenesis.

### Validation in human brain tissue

Single-nucleus RNA sequencing analysis of *C9orf72* patient cortex provides crucial validation of our in vitro findings. The identification of excitatory neuron subclusters with elevated G1/S and S-phase scores confirms that post-mitotic neurons inappropriately re-enter the cell cycle in human disease. The heterogeneous vulnerability across neuronal subclusters (with subclusters 32 and 35 most severely affected) may explain selective neuronal loss patterns in *C9orf72* ALS/FTD. Furthermore, our copy number variation analysis reveals genomic instability affecting DNA repair and cell cycle regulatory pathways, providing a potential mechanistic link between DNA damage and cell cycle dysregulation. The enrichment of alternative NHEJ repair genes in CNV gain regions and cell cycle transition genes in loss regions suggests a complex interplay between genome maintenance and cell cycle control that warrants further investigation.

## Conclusions and future directions

This work establishes cell cycle dysregulation as a central, druggable mechanism in *C9orf72* ALS/FTD. By demonstrating that arginine-containing DPRs drive CDK4/6 pathway activation, and that pharmacological inhibition rescues neuronal survival, we provide a potential path toward clinical translation. The availability of FDA-approved CDK4/6 inhibitors with CNS penetration enables immediate clinical trial design. Future studies should explore combination therapies targeting both cell cycle and DNA repair pathways, investigate the long-term effects of CDK4/6 inhibition in animal models, and determine optimal therapeutic windows for intervention. Additionally, the heterogeneous neuronal vulnerability we observe suggests that precision approaches targeting specific neuronal subtypes may maximize therapeutic benefit.

## Supporting information

Supplementary Figure 1. Relative mRNA expression of cell cycle regulation genes CCNC, CCND1, CCND2, CCNE1, CCNE2, CDK1, CDK2 and in iPSC derived C9or

## REFERENCES

1. DeJesus-Hernandez, M., et al. (2011). “Expanded GGGGCC hexanucleotide repeat in noncoding region of C9ORF72 causes chromosome 9p-linked FTD and ALS.” Neuron 72(2): 245–256.

2. Renton, A. E., et al. (2011). “A hexanucleotide repeat expansion in C9ORF72 is the cause of chromosome 9p21-linked ALS-FTD.” Neuron 72(2): 257–268.

3. Gao, F. B., et al. (2017). “Dysregulated molecular pathways in amyotrophic lateral sclerosis-frontotemporal dementia spectrum disorder.” EMBO J 36(20): 2931–2950.

4. Yang, D., et al. (2015). “FTD/ALS-associated poly(GR) protein impairs the Notch pathway and is recruited by poly(GA) into cytoplasmic inclusions.” Acta Neuropathol 130(4): 525–535.

5. Zhang, Y. J., et al. (2019). “Heterochromatin anomalies and double-stranded RNA accumulation underlie C9orf72 poly(PR) toxicity.” Science 363(6428).

6. Lopez-Gonzalez, R., et al. (2016). “Poly(GR) in C9ORF72-Related ALS/FTD Compromises Mitochondrial Function and Increases Oxidative Stress and DNA Damage in iPSC-Derived Motor Neurons.” Neuron 92(2): 383–391.

7. Farg, M. A., et al. (2017). “The DNA damage response (DDR) is induced by the C9orf72 repeat expansion in amyotrophic lateral sclerosis.” Hum Mol Genet 26(15): 2882–2896.

8. He, L., et al. (2022). “C9orf72 functions in the nucleus to regulate DNA damage repair.” Cell Death Differ.

9. Walker, C., et al. (2017). “C9orf72 expansion disrupts ATM-mediated chromosomal break repair.” Nat Neurosci 20(9): 1225–1235.

10. Robinson, H., et al. (2022). “Telomere Attrition in Induced Pluripotent Stem Cell-Derived Neurons From ALS/FTD-Related C9ORF72 Repeat Expansion Carriers.” Front Cell Dev Biol 10: 874323.

11. Lopez-Gonzalez, R., et al. (2019). “Partial inhibition of the overactivated Ku80-dependent DNA repair pathway rescues neurodegeneration in C9ORF72-ALS/FTD.” Proc Natl Acad Sci U S A 116(19): 9628–9633.

12. Maor-Nof, M., et al. (2021). “p53 is a central regulator driving neurodegeneration caused by C9orf72 poly(PR).” Cell 184(3): 689–708 e620.

13. Kruman, II, et al. (2004). “Cell cycle activation linked to neuronal cell death initiated by DNA damage.” Neuron 41(4): 549–561.

14. Joseph, C., et al. (2020). “Cell Cycle Deficits in Neurodegenerative Disorders: Uncovering Molecular Mechanisms to Drive Innovative Therapeutic Development.” Aging Dis 11(4): 946–966.

15. Park, D. et al. (1998). Cyclin-dependent kinases participate in death of neurons evoked by DNA-damaging agents. J Cell Biol. 143:457-467.

16. Nandakumar, S., et al. (2021). “Cell Cycle Re-entry in the Nervous System: From Polyploidy to Neurodegeneration.” Front Cell Dev Biol 9: 698661.

17. Barnum, K. J. and M.J. O’Connell (2014). “Cell cycle regulation by checkpoints.” Methods Mol Biol 1170: 29–40.

18. Vazquez-Villasenor, I., et al. (2021). “Persistent DNA damage alters the neuronal transcriptome suggesting cell cycle dysregulation and altered mitochondrial function.” Eur J Neurosci 54(9): 6987–7005.

19. Ranganathan, S. and R. Bowser (2010). “p53 and Cell Cycle Proteins Participate in Spinal Motor Neuron Cell Death in ALS.” Open Pathol J 4: 11–22.

20. Nguyen, M. D., et al. (2001). “Deregulation of Cdk5 in a mouse model of ALS: toxicity alleviated by perikaryal neurofilament inclusions.” Neuron 30(1): 135–147.

21. Vazquez-Villasenor, I., et al. (2020). “Expression of p16 and p21 in the frontal association cortex of ALS/MND brains suggests neuronal cell cycle dysregulation and astrocyte senescence in early stages of the disease.” Neuropathol Appl Neurobiol 46(2): 171–185.

22. Zhang, Z., et al. (2013). “Downregulation of microRNA-9 in iPSC-derived neurons of FTD/ALS patients with TDP-43 mutations.” PLoS One 8(10): e76055.

23. Freibaum, B. D., et al. (2015). “GGGGCC repeat expansion in C9orf72 compromises nucleocytoplasmic transport.” Nature 525(7567): 129–133.

24. Wu, D., et al. (2024). “Neuronal cell cycle reentry events in the aging brain are more prevalent in neurodegeneration and lead to cellular senescence.” PLoS Biol 22(4): e3002559.

25. Sakae, N., et al. (2018). “Poly-GR dipeptide repeat polymers correlate with neurodegeneration and Clinicopathological subtypes in C9ORF72-related brain disease.” Acta Neuropathol Commun 6(1): 63.

26. Wang, H., et al. (2021). “DNA Damage and Repair Deficiency in ALS/FTD-Associated Neurodegeneration: From Molecular Mechanisms to Therapeutic Implication.” Front Mol Neurosci 14: 784361.

27. Liu, Q., et al. (2020). “To control or to be controlled? Dual roles of CDK2 in DNA damage and DNA damage response.” DNA Repair (Amst) 85: 102702.

28. McGurk, L., et al. (2018). “Nuclear poly(ADP-ribose) activity is a therapeutic target in amyotrophic lateral sclerosis.” Acta Neuropathol Commun 6(1): 84.

