## Supplementary Figure 1. Relative mRNA expression of cell cycle regulation genes CCNC, CCND1, CCND2, CCNE1, CCNE2, CDK1, CDK2 and in iPSC derived C9or for "Cell cycle dysregulation contributes to neurodegeneration in human neurons and defines a druggable vulnerability in *C9orf72* ALS/FTD"

### Slide 1
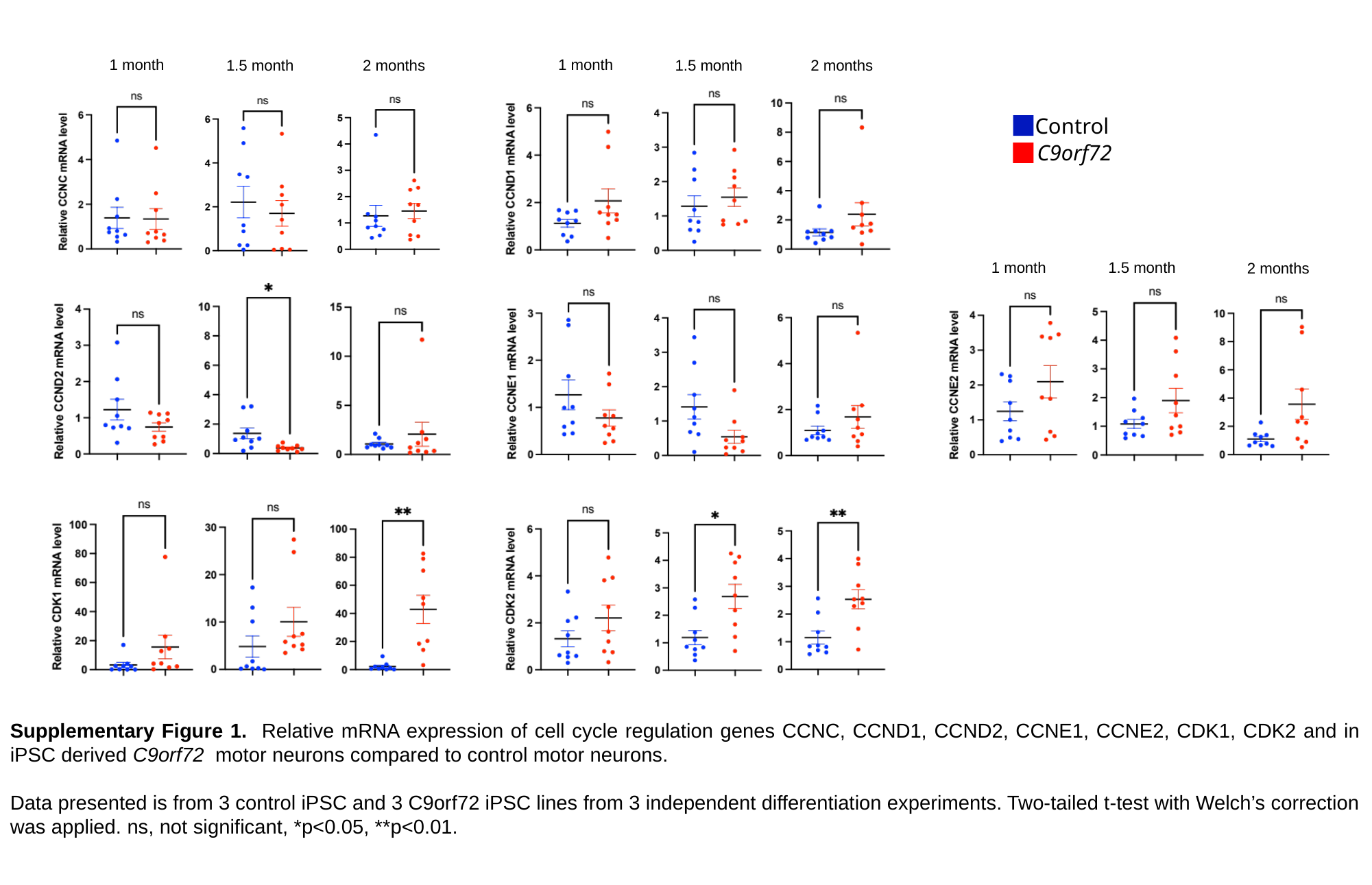

1 month
1 month
1.5 month
1.5 month
2 months
2 months
Control
C9orf72
1 month
1.5 month
2 months
Supplementary Figure 1. Relative mRNA expression of cell cycle regulation genes CCNC, CCND1, CCND2, CCNE1, CCNE2, CDK1, CDK2 and in iPSC derived C9orf72 motor neurons compared to control motor neurons.
Data presented is from 3 control iPSC and 3 C9orf72 iPSC lines from 3 independent differentiation experiments. Two-tailed t-test with Welch’s correction was applied. ns, not significant, *p<0.05, **p<0.01.

### Slide 2
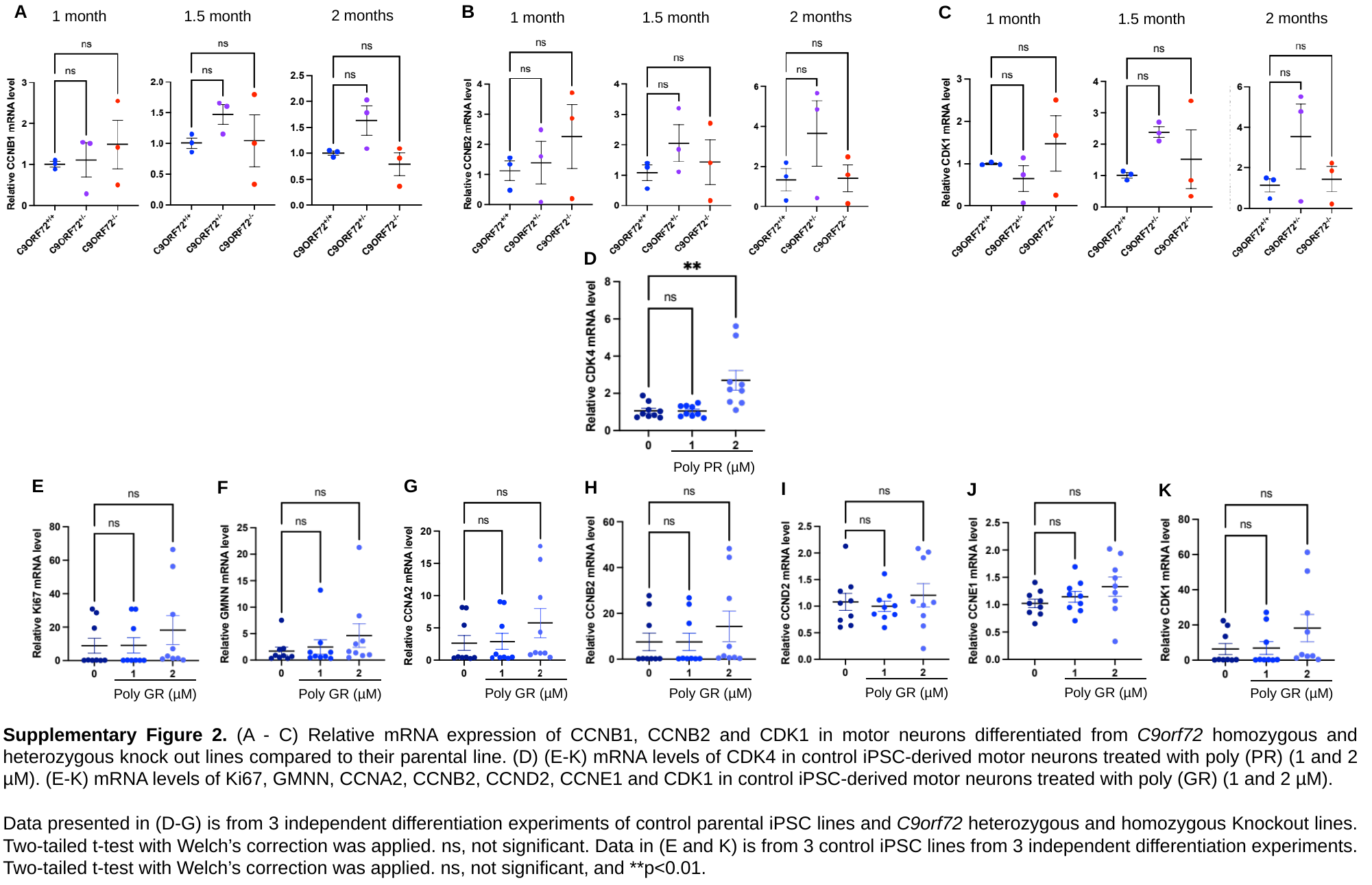

B
A
C
2 months
1.5 month
1 month
2 months
1.5 month
1 month
2 months
1.5 month
1 month
D
Poly PR (µM)
G
E
F
H
I
J
K
Poly GR (µM)
Poly GR (µM)
Poly GR (µM)
Poly GR (µM)
Poly GR (µM)
Poly GR (µM)
Poly GR (µM)
Supplementary Figure 2. (A - C) Relative mRNA expression of CCNB1, CCNB2 and CDK1 in motor neurons differentiated from C9orf72 homozygous and heterozygous knock out lines compared to their parental line. (D) (E-K) mRNA levels of CDK4 in control iPSC-derived motor neurons treated with poly (PR) (1 and 2 µM). (E-K) mRNA levels of Ki67, GMNN, CCNA2, CCNB2, CCND2, CCNE1 and CDK1 in control iPSC-derived motor neurons treated with poly (GR) (1 and 2 µM).
Data presented in (D-G) is from 3 independent differentiation experiments of control parental iPSC lines and C9orf72 heterozygous and homozygous Knockout lines. Two-tailed t-test with Welch’s correction was applied. ns, not significant. Data in (E and K) is from 3 control iPSC lines from 3 independent differentiation experiments. Two-tailed t-test with Welch’s correction was applied. ns, not significant, and **p<0.01.

### Slide 3
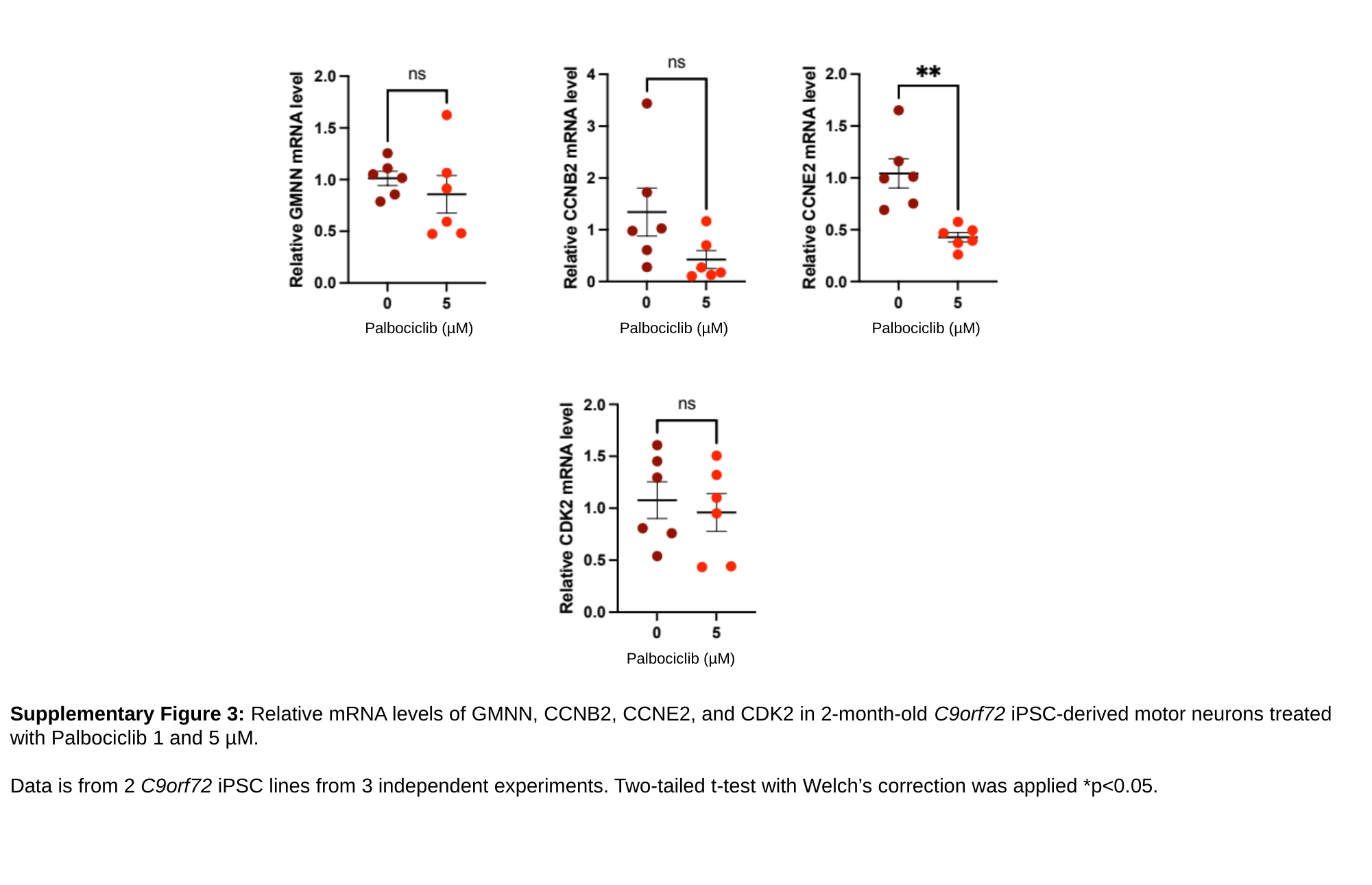

Palbociclib (µM)
Palbociclib (µM)
Palbociclib (µM)
Palbociclib (µM)
Supplementary Figure 3: Relative mRNA levels of GMNN, CCNB2, CCNE2, and CDK2 in 2-month-old C9orf72 iPSC-derived motor neurons treated with Palbociclib 1 and 5 µM.
Data is from 2 C9orf72 iPSC lines from 3 independent experiments. Two-tailed t-test with Welch’s correction was applied *p<0.05.

### Slide 4
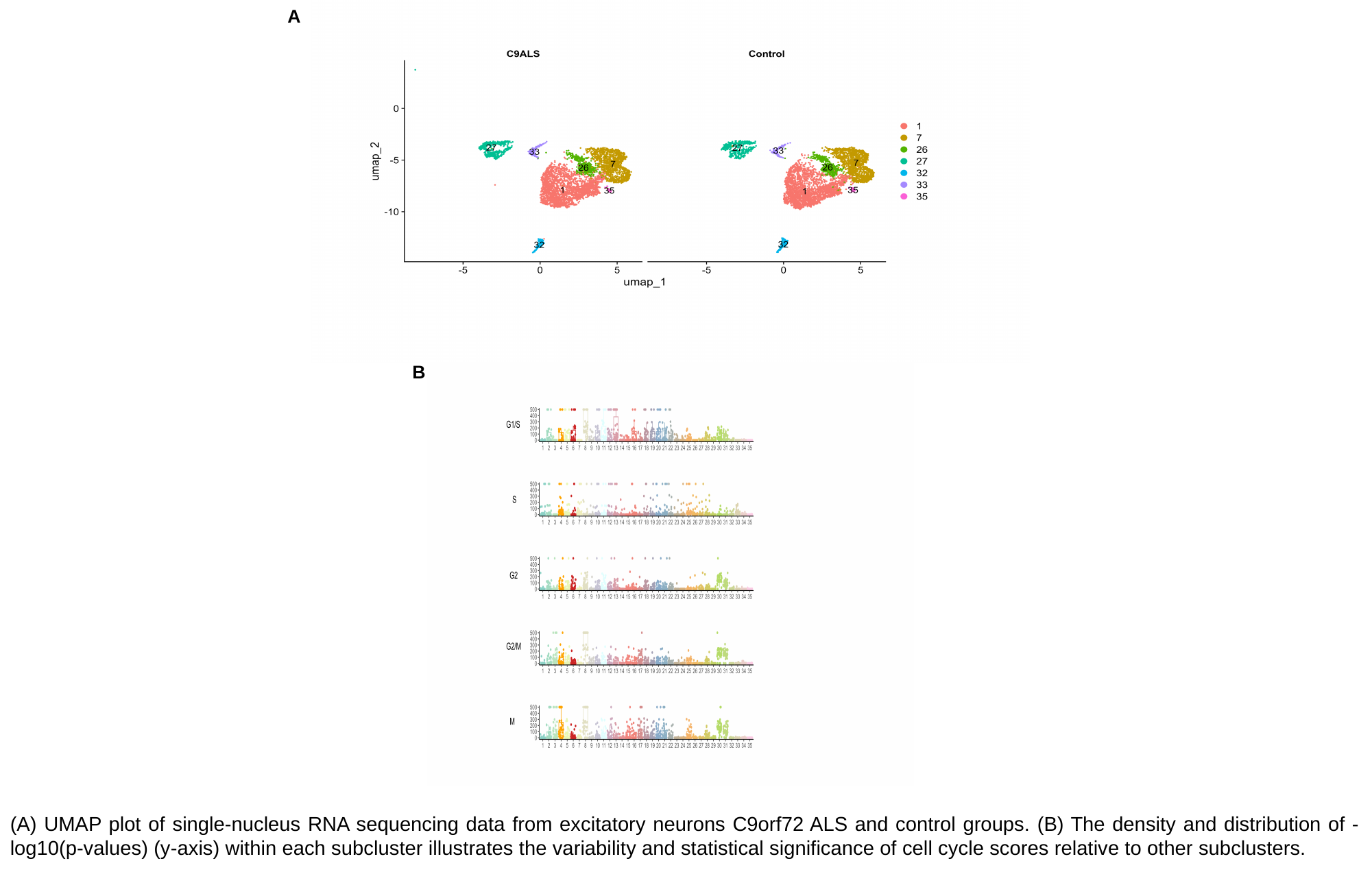

A
B
(A) UMAP plot of single-nucleus RNA sequencing data from excitatory neurons C9orf72 ALS and control groups. (B) The density and distribution of -log10(p-values) (y-axis) within each subcluster illustrates the variability and statistical significance of cell cycle scores relative to other subclusters.

### Slide 5
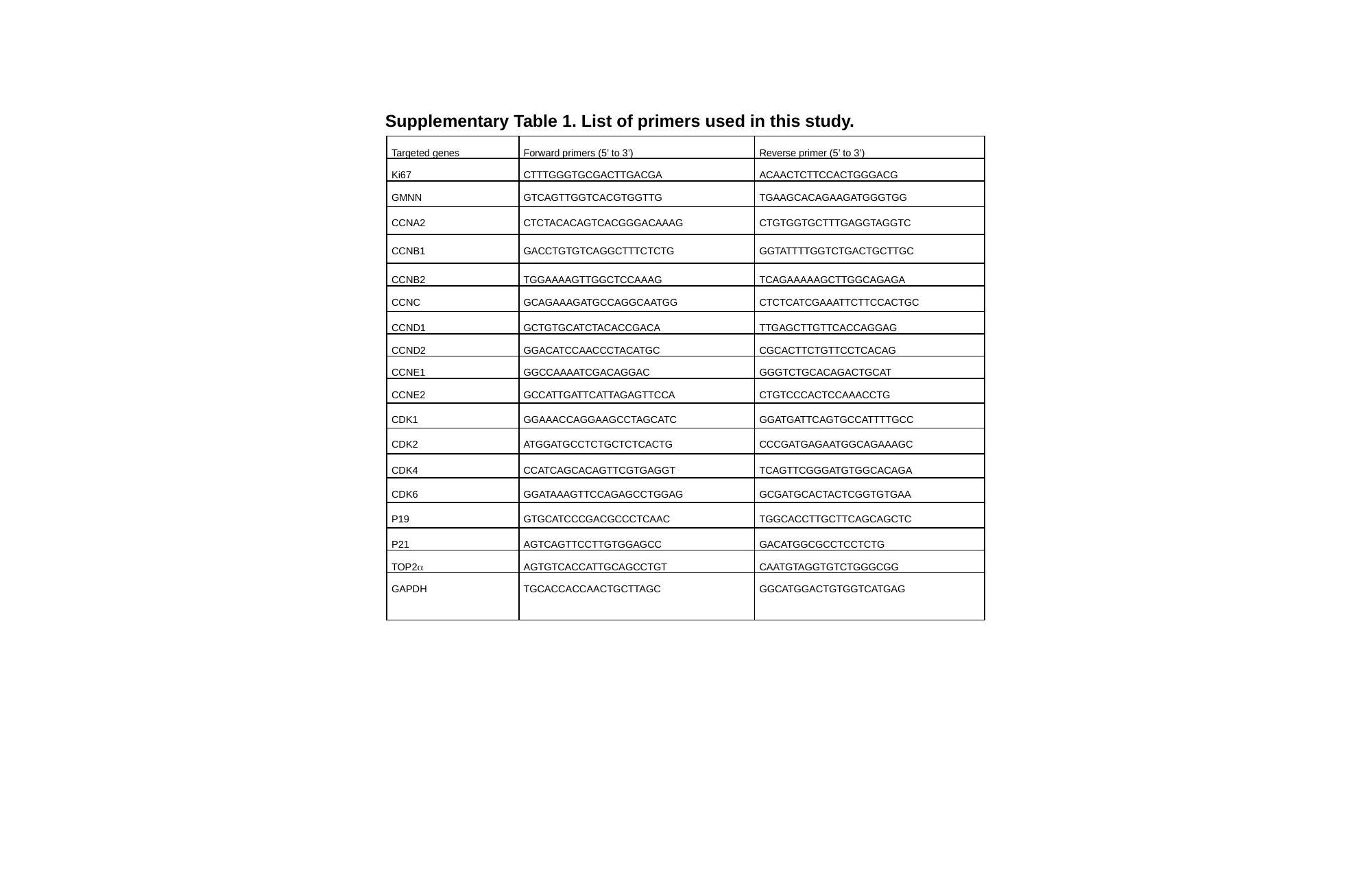

Supplementary Table 1. List of primers used in this study.
| Targeted genes | Forward primers (5’ to 3’) | Reverse primer (5’ to 3’) |
| --- | --- | --- |
| Ki67 | CTTTGGGTGCGACTTGACGA | ACAACTCTTCCACTGGGACG |
| GMNN | GTCAGTTGGTCACGTGGTTG | TGAAGCACAGAAGATGGGTGG |
| CCNA2 | CTCTACACAGTCACGGGACAAAG | CTGTGGTGCTTTGAGGTAGGTC |
| CCNB1 | GACCTGTGTCAGGCTTTCTCTG | GGTATTTTGGTCTGACTGCTTGC |
| CCNB2 | TGGAAAAGTTGGCTCCAAAG | TCAGAAAAAGCTTGGCAGAGA |
| CCNC | GCAGAAAGATGCCAGGCAATGG | CTCTCATCGAAATTCTTCCACTGC |
| CCND1 | GCTGTGCATCTACACCGACA | TTGAGCTTGTTCACCAGGAG |
| CCND2 | GGACATCCAACCCTACATGC | CGCACTTCTGTTCCTCACAG |
| CCNE1 | GGCCAAAATCGACAGGAC | GGGTCTGCACAGACTGCAT |
| CCNE2 | GCCATTGATTCATTAGAGTTCCA | CTGTCCCACTCCAAACCTG |
| CDK1 | GGAAACCAGGAAGCCTAGCATC | GGATGATTCAGTGCCATTTTGCC |
| CDK2 | ATGGATGCCTCTGCTCTCACTG | CCCGATGAGAATGGCAGAAAGC |
| CDK4 | CCATCAGCACAGTTCGTGAGGT | TCAGTTCGGGATGTGGCACAGA |
| CDK6 | GGATAAAGTTCCAGAGCCTGGAG | GCGATGCACTACTCGGTGTGAA |
| P19 | GTGCATCCCGACGCCCTCAAC | TGGCACCTTGCTTCAGCAGCTC |
| P21 | AGTCAGTTCCTTGTGGAGCC | GACATGGCGCCTCCTCTG |
| TOP2α | AGTGTCACCATTGCAGCCTGT | CAATGTAGGTGTCTGGGCGG |
| GAPDH | TGCACCACCAACTGCTTAGC | GGCATGGACTGTGGTCATGAG |

### Slide 6
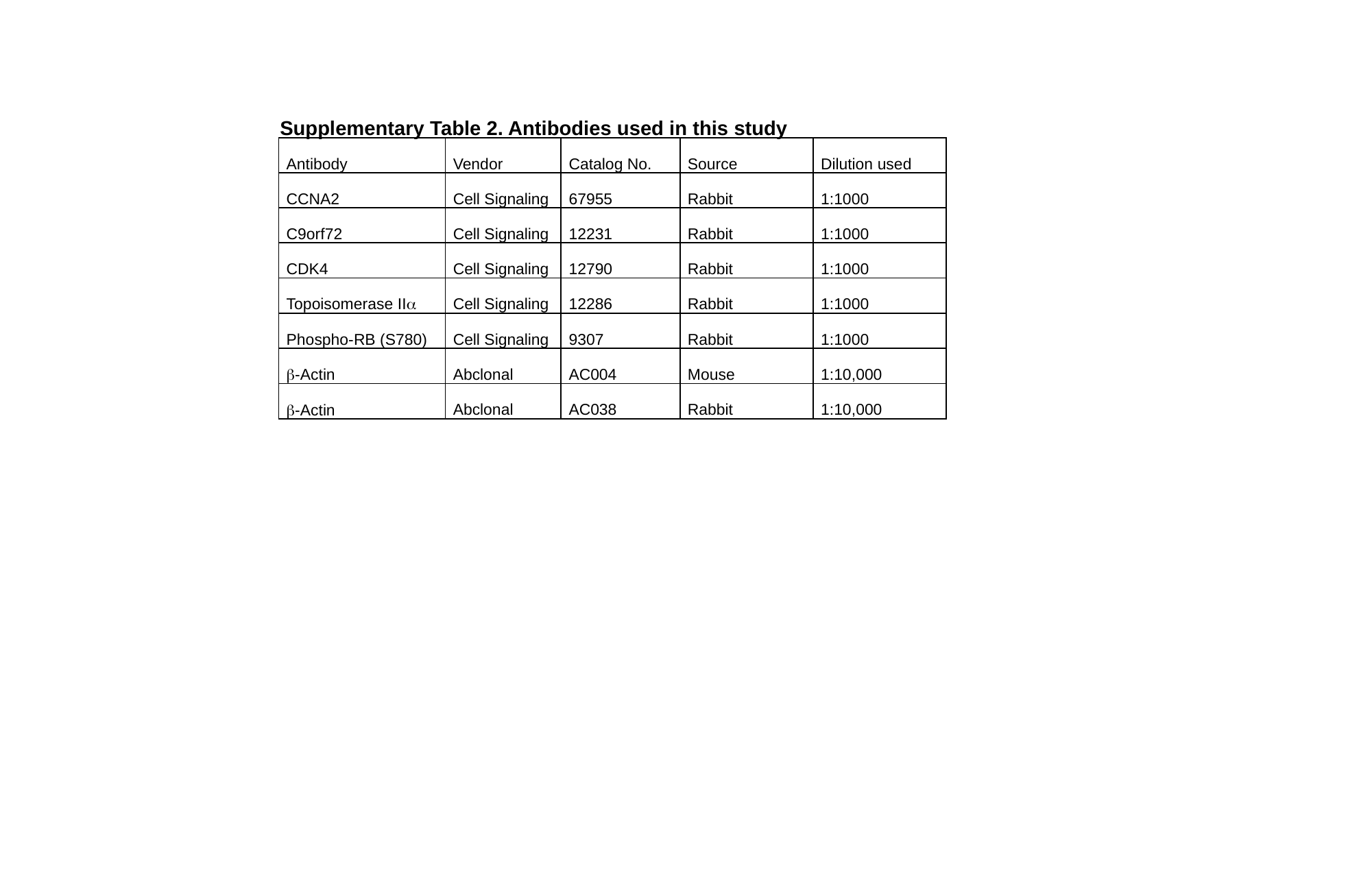

Supplementary Table 2. Antibodies used in this study
| Antibody | Vendor | Catalog No. | Source | Dilution used |
| --- | --- | --- | --- | --- |
| CCNA2 | Cell Signaling | 67955 | Rabbit | 1:1000 |
| C9orf72 | Cell Signaling | 12231 | Rabbit | 1:1000 |
| CDK4 | Cell Signaling | 12790 | Rabbit | 1:1000 |
| Topoisomerase IIα | Cell Signaling | 12286 | Rabbit | 1:1000 |
| Phospho-RB (S780) | Cell Signaling | 9307 | Rabbit | 1:1000 |
| β-Actin | Abclonal | AC004 | Mouse | 1:10,000 |
| β-Actin | Abclonal | AC038 | Rabbit | 1:10,000 |
